# A Partially Phase-Separated Genome Sequence Assembly of the *Vitis* Rootstock ‘Börner’ (*Vitis riparia* x *Vitis cinerea*) and its Exploitation for Marker Development and Targeted Mapping

**DOI:** 10.1101/854687

**Authors:** Daniela Holtgräwe, Thomas Rosleff Sörensen, Ludger Hausmann, Boas Pucker, Prisca Viehöver, Reinhard Töpfer, Bernd Weisshaar

**Affiliations:** Faculty of Biology, Center for Biotechnology, Bielefeld University, Bielefeld, Germany; Institute for Grapevine Breeding Geilweilerhof, Julius Kuehn-Institute, Federal Research Centre for Cultivated Plants, Siebeldingen, Germany

**Keywords:** de novo genome assembly, Vitis, rootstock, targeted mapping, SSR markers, SNV detection, whole genome sequencing

## Abstract

Grapevine breeding becomes highly relevant due to upcoming challenges like climate change, a decrease in the number of available fungicides, increasing public concern about plant protection, and the demand for a sustainable production. Downy mildew caused by *Plasmopara viticola* is one of the most devastating diseases worldwide of cultivated *Vitis vinifera*. Therefore, in modern breeding programs genetic marker technologies and genomic data are used to develop new cultivars with defined and stacked resistance loci. Potential sources of resistance are wild species of American or Asian origin. The interspecific hybrid of *Vitis riparia* Gm 183 x *V. cinerea* Arnold, available as the rootstock cultivar ‘Börner’, carries several relevant resistance loci. We applied next generation sequencing to enable the reliable identification of simple sequence repeats (SSR) and also generated a draft genome sequence assembly of ‘Börner’ to access genome wide sequence variations in a comprehensive and highly reliable way. These data were used to cover the ‘Börner’ genome with genetic marker positions. A subset of these marker positions was used for targeted mapping of the *P. viticola* resistance locus, *Rpv14*, to validate the marker position list. Based on the reference genome sequence PN40024, the position of this resistance locus can be narrowed down to less than 0.5 Mbp on chromosome 5.

## 1 Introduction

Downy mildew of grapevine is caused by the oomycete *Plasmopara viticola*. It is a serious threat for viticulture especially in humid and warm environments. Intensive chemical plant protection is necessary to protect grapevines under disease promoting conditions - frequently by using copper. Resistance breeding allows the selection of new cultivars that require a reduced number of chemical treatments. Genetic analyses of resistance loci promoted such breeding initiatives which today are oriented towards stacking (combining) of several resistance loci (Töpfer and Eibach, 2016). Therefore, in the recent past a number of resistance loci for downy mildew have been identified and were localized within grapevine genomes. Around two dozen loci designated *Rpv* (Resistance *Plasmopara viticola*) are listed in the Vitis International Variety Catalogue (VIVC, www.vivc.de, Hausmann et al., 2019)). Some of them are in elite genetic background like *Rpv1* (Merdinoglu et al., 2003), *Rpv3* (Welter et al., 2007; Bellin et al., 2009), *Rpv10* (Schwander et al., 2012), or *Rpv12* (Venuti et al., 2013) and can be used in breeding programs for selection of new cultivars. Other loci are less well characterized and essentially known only from wild species. To allow their application these loci need to be introgressed into favorable *Vitis vinifera* genetic background. Such an introgression can be accelerated considerably by using marker-assisted selection or marker-assisted back crossing (Herzog et al., 2013). To start such an approach a locus should be well characterized and sufficient closely linked markers should be available. Within plant breeding programs simple sequence repeats (SSRs) and single nucleotide variant (SNVs) markers are widely applied. The discovery and development of this marker types by using NGS data has been successfully demonstrated for various plant species (Taheri et al., 2018).

Since the grapevine reference genome sequence became available (Jaillon et al., 2007), QTL mapping by using SSR markers is simplified in a reference sequence guided approach (e.g. (Schwander et al., 2012)). Alternatively, next generation sequencing approaches (NGS) were successfully used to speed up mapping processes (Barba et al., 2014; Hyma et al., 2015; Capistrano-Gossmann et al., 2017).

One of the genetically useful genotypes as a source for resistances is the rootstock cultivar ‘Börner’. It is an interspecific hybrid of *V. riparia* Gm 183 × *V. cinerea* Arnold, so ‘Börner’ is derived from individuals of two *Vitis* species native to North America that carry several resistances. Among these are resistances for phylloxera (*Rdv1* - (Zhang et al., 2009)), black rot (*Rgb1* and *Rgb2* - (Rex et al., 2014)), and for downy mildew (*Rpv14* - (Ochssner et al., 2016)). To support molecular analysis of these and other loci we generated a first draft genome sequence assembly of the highly heterozygous ‘Börner’ genome. We produced comprehensive lists of SSR and SNV marker positions based on NGS data from ‘Börner’, validated these marker positions randomly as well as by using *Rpv14* on chromosome 5 (chr.5) as an example target.

The first grapevine reference genome sequence was generated based on Sanger sequencing technology using the ‘Pinot noir’ inbreeding line PN40024, which is estimated to be homozygous for 93% of the genome, thus enabling a successful assembly ((Jaillon et al., 2007) currently version 12×.v2). However, PN40024 suffers from inbreeding depression and selfing grapevine for homozygosity is difficult and a time consuming process. Other available grapevine genome sequences until 2018 were derived from heterozygous genotypes (Velasco et al., 2007; Adam-Blondon et al., 2011; Di Genova et al., 2014), posing a challenging sequencing and assembly problem which is not yet fully solved. In 2007, it was impossible to achieve a high quality genome assembly for ‘Pinot noir’ due to its heterozygosity since merged allelic sequences made it impossible to separate both haplophases. As NGS techniques developed progressively, the number of sequenced plant genomes has rapidly increased breaching the 100 mark some time ago (Michael and Jackson, 2013; VanBuren et al., 2015). Only a few genome sequences are originally based purely on the Sanger sequencing technique (e.g. *Arabidopsis* and grapevine). A larger number was generated based on mixed data types including Sanger, 454 and Illumina (e.g. potato, sugarbeet, *Brassica rapa*) or only Illumina derived sequences (e.g. eggplant, Norway spruce). The quality of the latter, generated by early NGS technologies, does not reach the quality of the older BAC-based Sanger assemblies despite higher read data coverage (Alkan et al., 2011; Burton et al., 2013). Although long reads became the best option for generating high quality genome sequences (Li et al., 2018), the high throughput and low costs of Illumina sequencing has made it economical to sequence plenty of genomes at least to a draft state or to re-sequence individual genotypes. A major problem and current challenge in plant genome sequence assembly is resolving the repeat structures and proper continuous separation of haplophases within heterozygous or polyploid plants (VanBuren et al., 2015). Repeat sequences are very common in plant genomes including repeats that are more than 10 kb in length. Therefore, resolving plant genome complexity with short NGS reads, mainly 100-300 nt in length, is a big challenge. Such assemblies result in collapsed sequences and break down at the borders of repetitive regions causing shortening of contigs. Some improvements were reached by using a variation of paired end libraries (VanBuren et al., 2015) combined with mate pair libraries. Long insert libraries were of particular value to bridge such gaps up to about 12kb.

Here we utilized sequencing data mainly generated by 454 and Illumina technologies to generate a draft assembly of the ‘Börner’ genome sequence with the main goal to allow easy access to markers. The approach was complemented by sequencing BAC ends from a library created from ‘Börner’ DNA. We hypothesized that the interspecific nature of ‘Börner’ and the genetic distance of its two wild ancestor species, *V. riparia* and *V. cinerea*, could be helpful to dissect and assign the haplophases of ‘Börner’ in its parental genomes even with Illumina reads. Having at least to a certain extent phased contigs at hand, several downstream applications become feasible. The alignment of ‘Börner’ contigs to the reference PN40024 sequence revealed many polymorphic sites including those at short sequence repeat (SSR) positions, being a source of potential high value for e.g. grapevine resistance breeding. The detection of single SNVs (single nucleotide variants) or groups of nearby SNVs within a multiple alignment of two ‘Börner’ contigs and the PN40024 reference sequence are considered to be useful in bulked segregant analysis (BSA). The identification of markers linked to loci or genes causing a disease resistance by applying BSA has been shown in plants (Michelmore et al., 1991). Pooling equal amounts of DNA is a practical way to reduce the cost of large-scale association studies to identify susceptibility loci for e.g. diseases (Sham et al., 2002). The pooling allows to measure allele frequencies in groups of individuals by using far fewer genotyping assays than are required if genotyping all individuals. Here we demonstrate the power of heterozygous positions within one species to physical narrow down a candidate locus for downy mildew resistance (*Rpv14*).

## 2 Material and Methods

### BAC library

To construct a BAC library of the grapevine cv. ‘Börner’, young frozen leaf material was sent to Bio S&T Inc. (Montreal, Canada) as a service provider for generation of BAC libraries. The DNA was inserted into the vector pIndigoBAC-5 (Epicentre, Madison, USA; ENA/Genbank accession number EU140754) cut with *Hin*dIII. Competent cells of *Escherichia coli* strain DH10B were transformed with the ligation mix and plated on LB medium supplemented with 12.5 μg/mL chloramphenicol (CM), 40 μg/mL X-GAL and 0.2 mM IPTG. White colonies were picked and transferred in 384-well microtiter plates filled with LB freezing medium (Zimmer and Verrinder Gibbins, 1997) containing CM. Altogether, 159 microtiter plates with 384 wells were generated which were duplicated and stored at −80°C. The insert size was estimated by pulsed-field gel electrophoresis to be on average 93 kbp. The library coverage is almost 12 haploid genome equivalents based on a genome size of about 500 Mbp (Lodhi and Reisch, 1995).

### BAC end sequencing by Sanger sequencing

A set of 43,008 (112 × 384) clones, comprising 8.4 haploid genome equivalents, of the ‘Börner’ BAC library (IGS-SCF-1083) was cultivated, harvested and applied to DNA extraction and subsequently sequenced from both ends on an ABI3730XL DNA analyzer (Applied Biosystems, Foster City, California, USA) as described before (Dohm et al., 2012). Sequencing primers were ccfw and ccrv for plate 1-52 and 55. Plates 53, 54 and 56-112 were sequenced using ccrv and a modified sequencing primer, pIfw (Supplementary Table 1).

### Processing of BAC end sequences (BES)

Trace files were processed by the base calling program phred (www.phrap.org, version 0.071220.c). Vector sequences were trimmed using the program blastall (Altschul et al., 1990) in version 2.2.24, with blast parameters set to the ones suggested by NCBI (www.ncbi.nlm.nih.gov/tools/vecscreen/about/). Low quality sequences were masked by applying a sliding window approach (size: 10 bp, minimal average score > 13) and the longest unmasked subsequence was taken as high quality sequence if the length was ≥ 80 bp. Slippage reads were removed as described before (Dohm et al., 2012). The dataset was filtered for contamination with *E. coli* sequences (blastall matches with an e-value ≤ 1e−50). Afterwards, the sequences were singularized per BAC end. A total of 69,444 high quality ‘Börner’ BES were submitted to ENA/GenBank and can be accessed under accession numbers KG622866 - KG692309.

### NGS sequencing library preparation

‘Börner’ DNA extracted from young leaf material with a commercial kit (Qiagen) according to the instructions of the manufacturer was sequenced using 454 GS-FLX Titanium (Roche), IonTorrent PGM (life technologies) as well as Illumina GAIIx, MiSeq and HiSeq1500. The preparation of Illumina paired-end (PE) and single-end (SE) sequencing libraries was performed according to the Illumina TruSeq DNA Sample Preparation v2 Guide. DNA was fragmented by nebulization. After end repair and A-tailing, individual indexed adaptors were ligated to the DNA fragments allowing multiplexed sequencing. The adaptor-ligated fragments were size selected on a 2 % low melt agarose gel to a size of 300-600 bp. After enrichment PCR of fragments that carry adaptors on both ends the final libraries were quantified with PicoGreen. The average fragment size of each library was determined on a BioAnalyzer High Sensitivity DNA chip. Libraries were sequenced either 100 nt on a SE run using one lane of a GAIIx run (ENA/SRA accession numbers ERR2729619, ERR2729620), or sequenced 2 × 100 nt PE on GAIIx (ERR2729739, ERR2729740, ERR2729741, ERR2729743, ERR2729744) or HiSeq1500 (ERR2729742) sequencers. A mate pair library was constructed according to the Illumina MatePair Sample Preparation v2 Guide. DNA was fragmented to a size of 3-4 kb by using Hydroshear. Fragment ends were repaired and biotinylated. Afterwards, fragments of 3.5 kb size were selected by gel electrophoresis. The resulting molecules were circularized. Removal of linear fragments was achieved by DNA exonuclease treatment. The circular molecules were randomly fragmented by nebulization. The biotinylated fragments were purified using streptavidin-coated magnetic beads. The following steps were carried out as described for the PE library construction. Finally, the library was sequenced in paired-end modus using an Illumina MiSeq generating 2 × 36 nt PE reads (ERR2729745).

GS-FLX sequencing libraries (454) were prepared from 10 micrograms of DNA according to the GS-FLX-Titanium Rapid Library Preparation Method Manual. DNA was fragmented by nebulization. Purified DNA fragments were subjected to end repair and subsequently to adaptor ligation. Large DNA fragments were selected via AMPure Beads. After emPCR and enrichment of DNA carrying beads sequencing was performed on several runs on a Roche/FLX sequencing instrument using Titanium XLR70 sequencing kits (ERR2728011-ERR2718029). Mate-pair libraries (ERR2728288-ERR2728292) with varying insert length (multi-span PE reads, 3 kbp and 8 kbp) were prepared from 15-20 micrograms of genomic DNA according to GS-FLX-Titanium manual method and using the GS FLX Titanium Paired End Adaptors kit and sequenced on a Roche/FLX sequencer.

The library construction for IonTorrent PGM (life technologies) sequencing was performed according to the Ion XpressTM Plus gDNA and Amplicon Library Preparation guide for a 200 bp library. DNA was fragmented by nebulization. The ends were repaired and ligated to adapters. Size selection of DNA molecules was performed via E-Gel® SizeSelectTM Agarose Gel to an average insert size of 350 bp. After enrichment of the template carrying Ion Sphere particles on an Ion OneTouch, sequencing was performed using a 318 chip on an Ion Personal Genome MachineTM (PGMTM) system (ERR2729812).

### NGS reads pre-processing

Illumina reads were quality trimmed and adapter clipped using Trimmomatic (Bolger et al., 2014) with all available adapter sequences from Illumina. Reads from the 454- and PGM/Ion Torrent-sequencing platform were trimmed with the CLC Genomics Workbench using the trim points of the SFF-files. Reads shorter than 36 nt were discarded. After pre-processing 75.62 Gbp raw data (=90.9% of the raw bases) were available for *de novo* assembly.

### Assembly, post-processing and validation

A *de novo* hybrid assembly of the ‘Börner’ genome was generated with the CLC Genomics Workbench (v. 8.0) using the following parameters: length fraction 0.5 similarity fraction 0.8, word size 20 and bubble size 50. For the re-mapping of the reads to the contigs that the CLC assembler carries out, the filtering thresholds were optimized to a length fraction of 0.8 (default: 0.5) and a similarity fraction of 0.95 (default: 0.9). From 75.62 Gbp of read data, representing nearly 80-fold coverage for each of the two ‘Börner’ haplophases, 338,484 contigs with a total length of 652.17 Mbp were calculated. A total of 91,3 % of the reads could be mapped back to the assembly, indicating a high probability that the sequence data was integrated correctly into the contigs. This intermediate raw assembly was designated “version 0.8.1”.

The assembly metadata include a value for the number of reads that cover a given position and also averages of these values for each contig. The value, which is essentially the total length of aligned reads divided by the assembly size, is below referred to as “alignment depth” (AD) and was used to deduce if a given contig represents a single haplophase or contains merged sequence information from both haplophases.

To test the completeness of the assembly we mapped the 69,444 ‘Börner’ BES on version 0.8.1. Mapping was done with the program BLAT (Kent, 2002). The majority (43,008 BES = 61.9%) mapped with 99% identity or better and for a length fraction of 0.5. About 4.5% BES matched the end of contigs with more than 20 to 50% of their length, but their mapping was not unique to one contig within the assembly. A small proportion of these (733 BES) produced multiple matches, suggesting that these contain mainly repetitive sequences. To avoid connecting contigs from different phases, we refrained from scaffolding since we had no information about the phase from which the BACs/BES were derived.

The assembly was checked for contaminations by searching against a custom database with sequences of human, yeast, mould, several fungal and viral pathogens along with insects such as aphids and mites with blast+ (e-value <=1e−50) (Camacho et al., 2009). A BLAST (Altschul et al., 1990) run querying GenBank’s non-redundant protein database (nr) completed the filtering process by using the same cut off criteria. Contigs matching to contaminants were also characterized by very low AD (1 - 20×). The removal of PhiX as well as sequences resembling cloning vectors caused a reduction of 213,834 bp and a total removal of three contigs. Plastid sequences from *V. vinifera* were detected in 67 contigs and removed as well. For subsequent analyses, contigs smaller than 500 bp were discarded. Although nearly 30% of all initial contigs were rejected by this filter, the total assembly size was only reduced by about 5%. After size filtering, 238,193 contigs remained and were included in the final assembly which was designated BoeWGS1.0 (Biosample PRJEB28084). The results were evaluated for standard assembly statistics parameters (Supplementary Table 2).

The BoeWGS1.0 contigs were aligned to the reference genome sequence PN40024 12×v2 (https://urgi.versailles.inra.fr/download/vitis/12Xv2_grapevine_genome_assembly.fa.gz) with blast+ with the dust filter turned off and using an e-value cutoff of 1e-50 and a culling limit of 1. As a result of this analysis, each mapping contig from BoeWGS1.0 receives location information derived from PN40024. For quite some loci, there are exactly two BoeWGS1.0 contigs assigned to one locus in the reference sequence, indicating that the assembly separated the two ‘Börner’ haplophases. We refer to these contigs, which may contain allelic sequence information, as “contig pairs”.

BWA-MEM (Li, 2013) was applied to map all PE Illumina reads to the final assembly sequence using the −M option to mark short splits as secondary. The resulting mapping was subjected to inspection by REAPR (Hunt et al., 2013) to assess the assembly quality based on coverage, distance of PE reads, and orientation of PE reads.

BUSCO v3 (Simão et al., 2015) was deployed to analyze the recovery of the ‘embryophyta odb9’ benchmarking sequences in the assembly. This analysis was performed in the ‘genome’ mode using an e-value cutoff of 1e-3 and considering up to three candidate regions per BUSCO hit.

### Genome-wide variant detection

A strict mapping of 245,325,650 ‘Börner’ PE reads (2×100 nt) on the PN40024 12×.v2 reference sequence with an average coverage of 50× formed the basis for the variant detection. The mapping was done with CLC Genomics Workbench, the similarity fraction set to 0.95 and the length fraction to 0.9. Non-specific matches were discarded. Variants (single nucleotide variants (SNVs), insertions as well as deletions (InDels) and multi nucleotide variations MNVs) were detected with CLCs Probabilistic Variant Caller using default parameters. Only bi-allelic SNVs with a read coverage between 10 and 90 were kept for downstream analyses, whereas tri-allelic and poly-allelic variants, InDels and MNVs were ignored.

Bi-allelic SNVs were classified as homozygous variants if both ‘Börner’ haplotypes differ from the reference sequence (in these cases both ‘Börner’ haplophases contain the same nucleotide at the inspected position), whereas heterozygous bi-allelic SNVs are those where only one ‘Börner’ haplotype differs from the reference sequence. The complete SNV data is available at https://doi.org/10.4119/unibi/2938178 and https://doi.org/10.4119/unibi//2938180.

### Simple sequence repeat (SSR) detection

SSR detection was done with MISA (MIcroSAtellite identification tool; http://pgrc.ipk-gatersleben.de/misa/misa.html, (Thiel et al., 2003)) using default parameters. Possible SSR marker positions were evaluated by comparison of the candidate SSRs plus adjacent +/−200 bases from BoeWGS1.0 with the SSRs of the PN40024 12×.v2 reference sequence. Comparison was performed with blastn with dust filter turned off and 1e-50 as e-value cutoff. SSR candidates where a BoeWGS1.0 contig pair matched one coordinate in the reference sequence and differences did not exceed 40 bp (identity >=90%) were classified as useful for SSR marker development. The SSR candidates were named “SSR [chr#]_[contig-ID]” with a trailing number if more than one SSR candidate was detected for one contig pair.

### SSR marker validation

For the validation of heterozygous SSRs within the BoeWGS1.0 assembly and variants in relation to the reference sequence, 53 SSR candidate positions were randomly selected for SSR marker validation. The SSRs from different genomic regions were selected with a minimum of two nucleotides and of six contiguous repeat units in the reference sequence. Primer3 (Untergasser et al., 2012) was used to design the amplimers (two primers directing towards each other based on the source sequence) fitting to the BoeWGS1.0 contig sequences as well as to the reference sequence PN40024 (Supplementary Table 1). The expected amplicon size of the markers was set from 150 to 400 bp and the primer size from 18 to 27 nt. DNA from ‘Börner’ and V3125 (see below) was used to check the polymorphism of the SSR marker sequences by PCR and gel electrophoresis on a 3% agarose gel. A few PCR products with expected smaller sequence variations were applied to Sanger sequencing and analyzed at the bp level by multiple sequence alignments. Verified heterozygous markers either in ‘Börner’ and/or in relation to V3125 were analyzed by fragment analysis using DNA of the crossing parents (V3125 and ‘Börner’) as well as the parental lines of ‘Börner’ as described before (Ochssner et al., 2016)

### Locus-guided variant calling and marker generation

The BoeWGS1.0 contig mapping on the PN40024 12×.v2 reference sequence was evaluated for a genomic region on chr.5. The BLAST results from the post-processing analysis described above were used. Regions where a multiple alignment composed of exactly two BoeWGS1.0 contigs (i.e. contig pairs) and parts of chr.5 from the PN40024 12×.v2 reference sequence was built were considered as genome regions representing both haplophases, if the contig pairs selected showed the expected AD of about 75× (75±25).

Each contig pair was aligned with edialign (from the EMBOSS suite, version 6.2) using default parameters. Alignments without N-stretches and with lengths of at least 500 bp were subjected to SNV detection. The list of contig pairs used for SNV detection on chr.5 is available as Supplementary Table 3. The downstream analysis focused on grouped SNVs between BoeWGS1.0 contig pairs and which are bi-allelic after comparison with the PN40024 reference sequence.

### Amplimer design for the *Rpv14* locus

BoeWGS1.0 contig pairs on chr.5 were used for SNV-based marker development addressing the *Rpv14* locus. Amplimers were designed using primer3 (Untergasser et al., 2012) preferably for contig pair alignments with a rate between 0.4 and 1 SNVs per 100 bp (Supplementary Table 1 and Supplementary Table 3). We targeted at an expected amplicon size between 400 and 530 bp. Deduced amplimers were tested to be unique in BoeWGS1.0 and PN40024 12×.v2.

### Plant material, bulk set up, DNA extraction and PCR

A biparental F1 mapping population of the grapevine breeding line V3125 (‘Schiava Grossa’ × ‘Riesling’) and ‘Börner’ (*V. riparia* Geisenheim 183 × *V. cinerea* Arnold), cultivated in the field at the JKI Institute for Grapevine Breeding Geilweilerhof, Siebeldingen, Germany, has been phenotyped three times for downy mildew leaf resistance as described previously (Ochssner et al., 2016). For SNV genotyping by amplicon sequencing genomic DNA was extracted from the parents V3125 and ‘Börner’ and selected F1 genotypes of the mapping population. Building bulks of DNA from all the individual samples and typing one SNV marker at a time can save valuable template DNA and has been successfully utilized in microsatellite markers (Barcellos et al., 1997) and SNPs (Shaw et al., 1998). Two bulks with 10 individuals each were generated from the mapping population. The first (R) and second (S) bulks encompass resistant and susceptible genotypes, respectively. Genomic DNA was extracted from young leaf material with a commercial kits (Qiagen) according to the instructions of the manufacturer. DNA samples were quantified using the Qubit™ dsDNA BR Assay Kit with the Qubit® 2.0 Fluorometer (Invitrogen, Life Technologies, Darmstadt, Germany).

Genomic DNA from the parents of the mapping population V3125 × ‘Börner’, as well as from the two bulks, was used as template in 25 μl PCR reactions with the two marker-specific primers and 1 ng DNA per individual under standard conditions. The amplicons obtained were purified and sequenced from both directions using the ABI Prism BigDye Terminator chemistry on an ABI Prism 3730 sequencer (Applied Biosystems).

### Determination of SNV/allele ratios

For the estimation of allele ratios only bi-allelic BoeWGS1.0 SNVs were considered that differentiate the two ‘Börner’ phases, i.e SNVs between BoeWGS1.0 contig pairs. Since only bi-allelic SNVs were considered, these positions are often homozygous in the V3125 genotype for one of the two ‘Börner’ alleles. Exceptions from this criteria (V3125 displays identical SNV heterozygosity to ‘Börner’) were excluded from the analysis. As a result, the allele frequency contributed from the ‘Börner’ parent has to be 50%, and is 100% for the V3125 parent when counting the variant that is present in V3125. Following these assumptions, the expected frequency for each single allele in a bulk of the unselected F1 individuals of this mapping population is 75%. We have estimated the SNV frequencies within the parental genotypes in order to confirm the determination, and to gain hints for problematic amplimers that amplify the haplophases with a bias for one of the two.

The estimation of the SNV frequencies among the pooled DNA samples (Pool R = resistant; Pool S = susceptible) and comparison with the parental lines was carried out by using the tool QVSanalyzer (Carr et al., 2009). The Sanger sequence trace files for determination of the relative proportions of two sequence variants were analyzed per batch. The generated output files contain details of the examined sequence variant ratios for individual samples, as well as summary statistics. The area below the peak at the position of the targeted SNV was calculated and set into relation to the surrounding peaks for each sample and corresponding trace file. For graphical presentation of the results, the ratios were converted into percent values for SNV or allele frequency. SNVs considered in the analysis are listed in Supplementary Table 3.

## 3 Results

### A draft assembly of the ‘Börner’ genome sequence

The data for the ‘Börner’ genome sequence were obtained by whole genome shotgun (WGS) sequencing. The assembly, designated BoeWGS1.0, has a total size of 618.3 Mbp, the N50 sequence size was 4,255 bp (Supplementary Table 2). We evaluated the average AD for the contigs in the assembly and classified the contigs with respect to the expected depth of aligned reads for paired contigs (separated haplophases) and contigs with merged sequences (both haplophases combined into one contig, see Supplementary Figure 1). A significant proportion of the contigs showed an AD between 50 and 100, that is 75±25 which can be considered as the expected value for a haplophase-specific contig. The range has been deduced from the total amount of reads (75 Gbp), 2 × 500 Mbp for the expected fully diploid sequence length and an interval of ±33%.

REAPR flagged a total of 14,007 regions as erroneous due to various reasons. Fragment coverage distribution errors within a contig account for 4,250 cases, while the same error type over a gap adds 5,674 cases. The remaining cases are contributed by low coverage within a contig or across a gap with 3,723 and 360 cases, respectively. In summary, the assessment by REAPR indicates a short read assembly from heterozygous material of acceptable to good quality.

BUSCO, i.e. the Benchmarking of Universal Single-Copy Orthologs, revealed the presence of 55.9% of all 1440 benchmarking genes from the reference set for embryophyta, 42% of the total 1440 as single copy and 13.3% as duplicated. In addition, fragments of 19.1% benchmarking genes were detected. Only 25% of the benchmarking genes were not detected in the assembly. It should be noted that the duplicated BUSCO genes can be explained by detection of two allelic versions in BoeWGS1.0 contig pairs.

### BoeWGS1.0 contigs representing different haplophases

The BoeWGS1.0 contigs were mapped against the *V. vinifera* reference genome sequence PN40024 12×.v2. About 210,444 (88.3%) could be mapped successfully with at least 30% of their length (https://doi.org/10.4119/unibi/2938185). The ratio of the total length of the mapped BoeWGS1.0 contigs to the length of the reference chromosomes is 1,14 on average (1.02 to 1.31; Supplementary Table 4), indicating uniform assembly quality and an overall homogeneous synteny between ‘Börner’ and PN40024. The median of the average contig ADs for all 19 chromosome mappings was 86.3 (Supplementary Table 4). Conspicuously, it is only the genetically unassigned chromosome “Ukn” or „unknown”, where the mapped BoeWGS1.0 contigs have an AD of 248 on average. The remaining, unmapped contigs contain sequences which are too diverse from the PN40024 reference sequence to be mapped.

More than one quarter (29.2%; 142 Mbp) of the PN40024 reference sequence was covered by BoeWGS1.0 contig pairs, again suggesting that the two ‘Börner’ haplophases were partially separated in the assembly. Most of these contigs show on average an AD in the expected range. Another quarter of the reference (28.7%; 139 Mbp) was covered by one BoeWGS1.0 contig, these contigs often (but not always) display higher AD values. In these cases, either the two ‘Börner’ haplophases were merged into one contig during assembly, or only one of the two haplophases displays sufficient similarity to become mapped.

Due to repetitive sequences, a small fraction of the PN40024 reference sequence (3.7%; 18 Mbp) is matched by more than two contigs. Nearly 38.5% (187 Mbp) of the reference sequence (including 3% N-stretches) is not covered by any stringently matching BoeWGS1.0 contig (Figure1).

**Figure 1:**
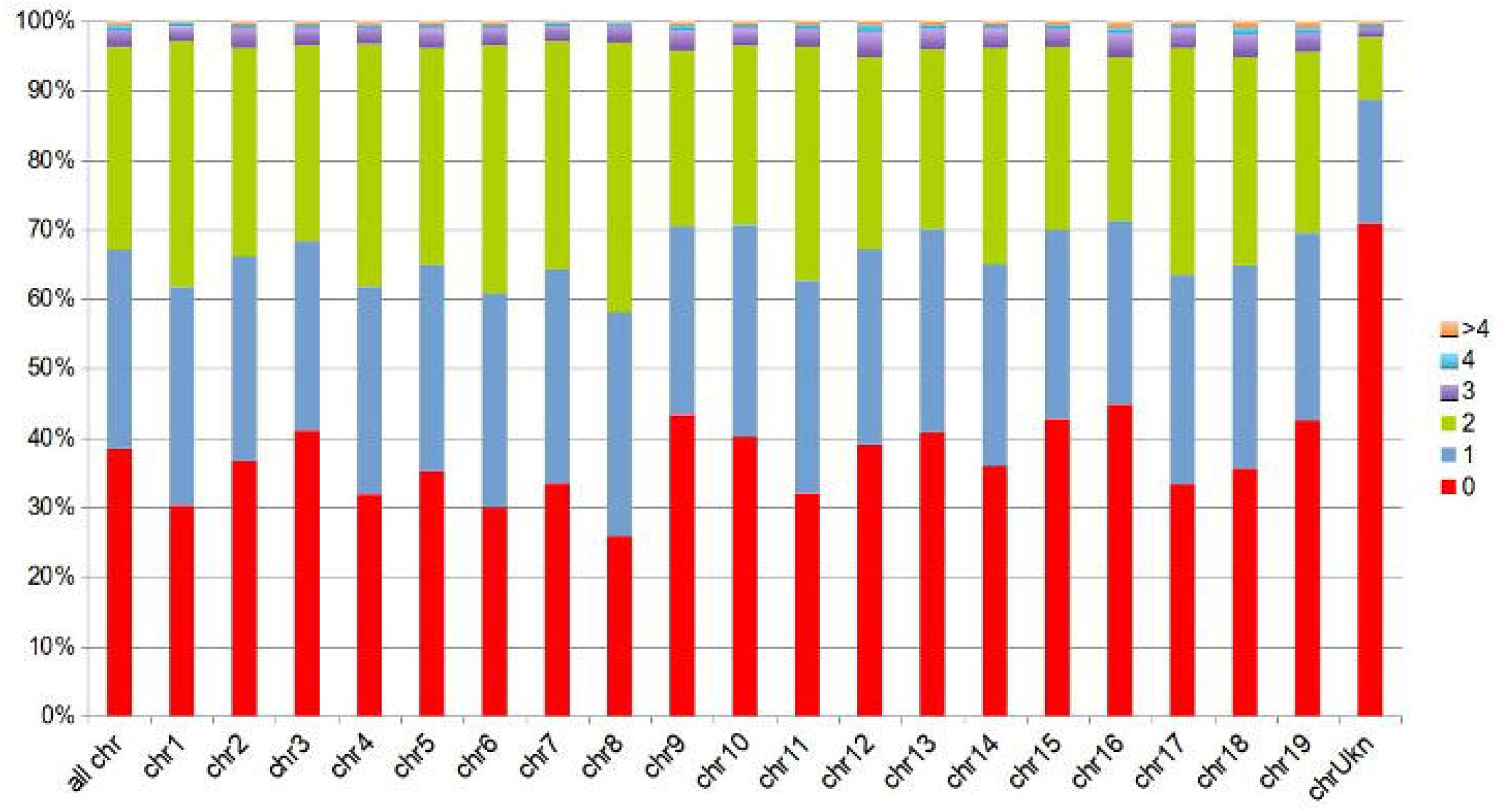
Sequence fractions of the PN40024 reference covered by BoeWGS1.0 contigs, displayed for all PN40024 pseudochromosomes individually (chr#), and for the whole PN40024 reference (allchr’s) in the leftmost column. Not covered fractions (given in percent) are shown in red (0), fractions covered by a single (1) BoeWGS1.0 contigs in blue, and fractions covered by contig pairs (2) in green. The remaining fractions are covered by three or more BoeWGS1.0 contigs (3, 4, >4).

### Homozygous and heterozygous variant frequencies

Genomic variants including SNVs contribute not only to intraspecific diversity but are candidates for valuable molecular markers. By mapping ‘Börner’ NGS reads against the reference sequence and subsequent variant calling, almost 5 million highly reliable bi-allelic SNVs were detected (https://doi.org/10.4119/unibi/2938178 and https://doi.org/10.4119/unibi/2938180)

Half of the 4,996,490 SNVs are heterozygous SNVs (2,536,406), which means one of the BoeWGS1.0 contigs shows the same nucleotide like the reference at a given position. The other half (2,460,084) are homozygous SNVs, describing variants only existing between PN40024 and BoeWGS1.0. The frequency is on average 1 per 77 bp (Supplementary Table 5), ranging from 1/70 to 1/82 bp for chr.3, chr.8 and chr.16 respectively. In addition to SNVs, MNVs and small InDels were called, dropping the overall variant frequency slightly to 1 variant per 68 bp.

### SSR detection for marker development

Simple sequence repeat (SSR) markers are often used to study molecular diversity or heterozygosity as well as for genetic mapping in grapevine. By comparing SSR positions in the grapevine reference genome sequence with BoeWGS1.0, a total of 10,820 putative SSR marker positions (“candidates”) with different unit sizes (two to six) and repeat numbers (up to 27) were deduced (Table1). Regions of the reference matched by a BoeWGS1.0 contig pair were exploited. Out of the 10,820 positions, 12% (1,313) were monomorph and 38% (4,110) were tri-allelic in an alignment of the three sequences. The remaining 50% were either bi-allelic or showed more than one motif in the SSR region. The more than 4,000 tri-allelic SSR candidates are the most valuable ones with regard to genetic mapping in a broad range of accessions (Supplementary Table 6). Not all of them are new because e.g. candidate SSR chr1_1203 corresponds to the established marker GF01-03, SSR chr1_1083 to marker GF01-21, and SSR chr1_1248 to marker GF01-22 (Fechter et al., 2014). However, the identification of already existing markers already indicates that our candidate list is valid.

**Table 1:**
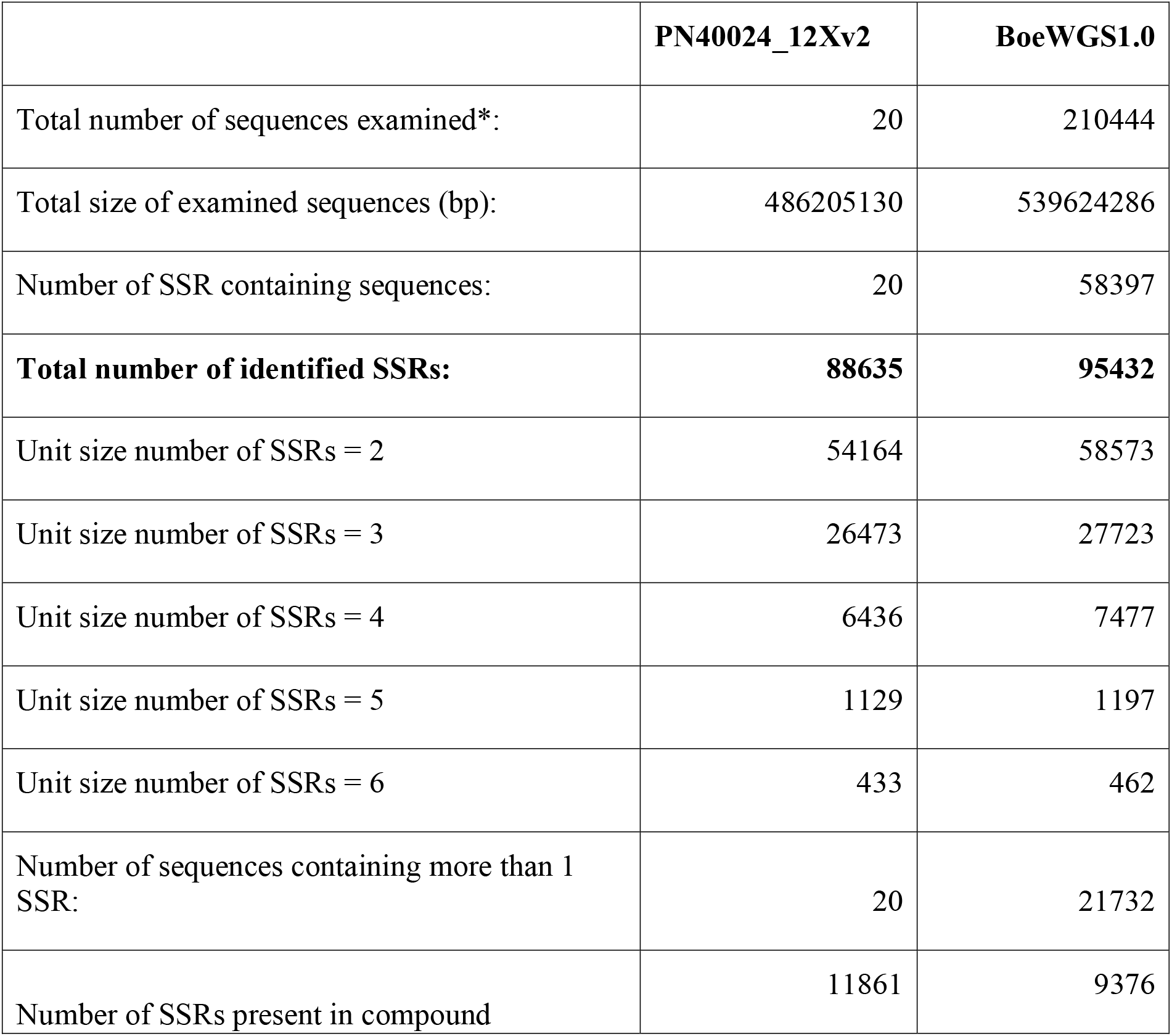
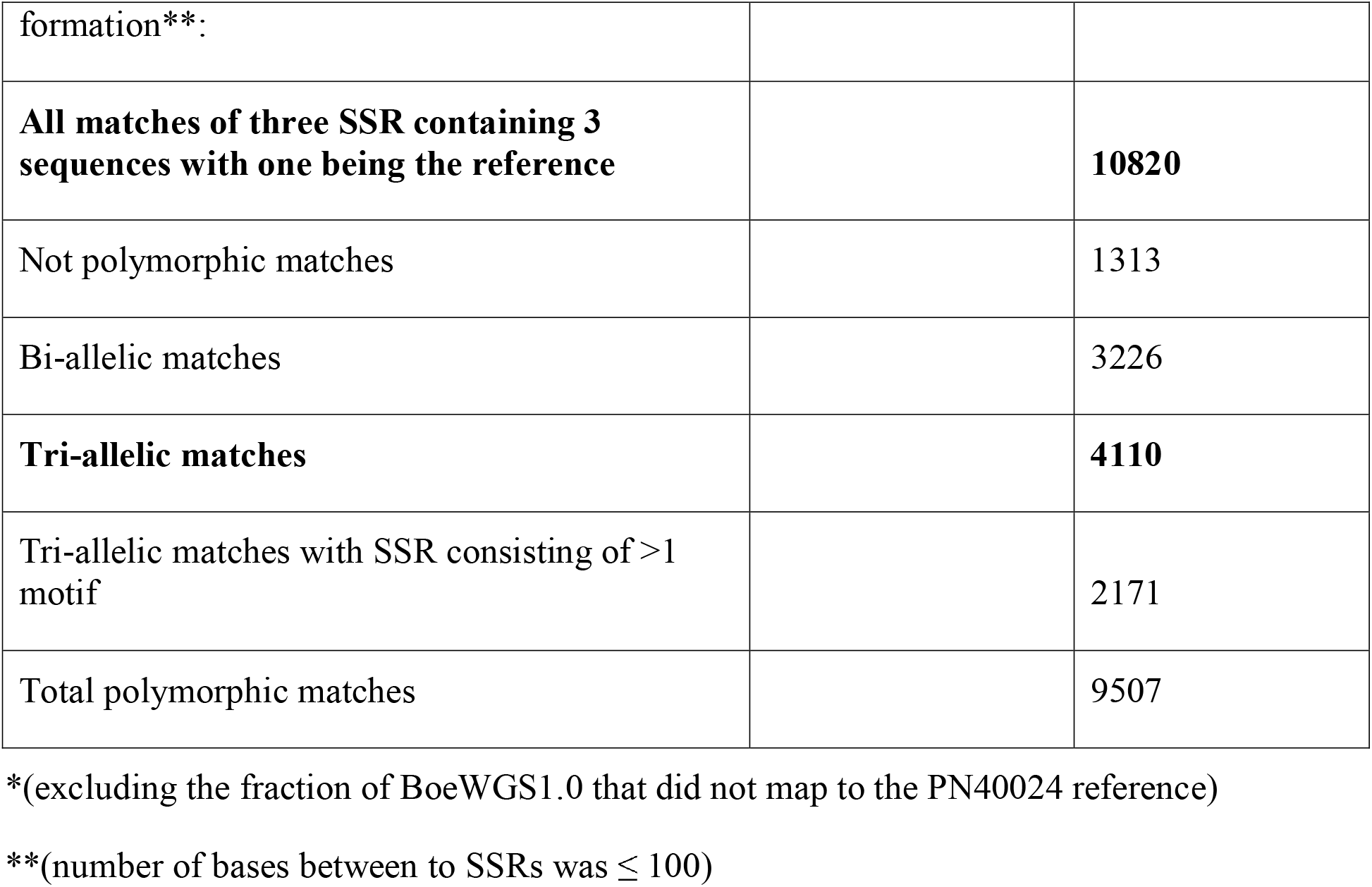
Mining and statistics of SSR containing sequences for marker development

|  | **PN40024_12Xv2** | **BoeWGS1.0** |
| --- | --- | --- |
| Total number of sequences examined*: | 20 | 210444 |
| Total size of examined sequences (bp): | 486205130 | 539624286 |
| Number of SSR containing sequences: | 20 | 58397 |
| **Total number of identified SSRs:** | **88635** | **95432** |
| Unit size number of SSRs = 2 | 54164 | 58573 |
| Unit size number of SSRs = 3 | 26473 | 27723 |
| Unit size number of SSRs = 4 | 6436 | 7477 |
| Unit size number of SSRs = 5 | 1129 | 1197 |
| Unit size number of SSRs = 6 | 433 | 462 |
| Number of sequences containing more than 1 SSR: | 20 | 21732 |
| Number of SSRs present in compound | 11861 | 9376 |
| formation**: |  |  |
| **All matches of three SSR containing 3 sequences with one being the reference** |  | **10820** |
| Not polymorphic matches |  | 1313 |
| Bi-allelic matches |  | 3226 |
| **Tri-allelic matches** |  | **4110** |
| Tri-allelic matches with SSR consisting of >1 motif |  | 2171 |
| Total polymorphic matches |  | 9507 |

\*(excluding the fraction of BoeWGS1.0 that did not map to the PN40024 reference)
\*\* (number of bases between to SSRs was $\leq 100$ )

A set of 45 tri-allelic and eight bi-allelic SSR candidates with a unit size of two or three randomly selected from all chromosomes was used to validate the power and reliability of SSR candidate prediction. The bi-allelic candidates were included to estimate the chance of false negative predictions. In this subset of eight candidates we tested six candidates, where the reference matches only one contig of BoeWGS1.0, and two candidates that were not found on a contig pair but share unit size and number. Out of the total set of 53 SSR sequences 38 could be verified by either PCR and subsequent agarose gel electrophoresis, Sanger sequencing or fragment analysis (Supplementary Table 7). The 15 negative cases were caused by unspecific primers, thus is the primers amplified various fragments instead of allelic amplicons. We did not optimize the amplimers empirically in a second round of amplimer design.

As expected ‘Börner’ amplicons with predicted small sequence variations between the alleles were analyzed more successful by Sanger sequencing than those with larger sequence variations. In six out of eigth cases the predicted bi-allelic SSR sequences were verified as monomorphic within ‘Börner’. Analysis of the sequenced nucleotides before and after the SSR region confirmed the assumption that these genome regions are less heterozygous than others (or monomorphic) and therefore only represented by a single contig in the BoeWGS1.0 assembly. For the SSR chr1_43023 candidate the new marker GF01-59 was established, although for this candidate only one BoeWGS1.0 contig matched to the reference. Both, Sanger sequencing and fragment analysis, confirmed the existence of two ‘Börner’ alleles for GF01-59.

Sanger sequencing of amplicons from SSR candidate loci with larger differences (e.g. in unit size, unit number, etc.) was less successful. In addition, we observed in these cases higher heterozygosity between the alleles even outside the SSR itself. However, validation by agarose gel electrophoresis worked out smoothly for these SSR candidates.

Since gel electrophoresis did not allow to verify exact allele sizes, these were determined by fragment analysis in 14 cases (Supplementary Table 7). Fragment analysis was performed with DNA of the exact parental lines of ‘Börner’ and V3125, in addition to ‘Börner’ itself. For 12 out of 14 SSR candidates different fragment sizes for the two ‘Börner’ alleles were detected that fit well to the predictions for both ‘Börner’ alleles. The validated SSR candidates for which we generated complete documentation received marker designations. Seven of these SSR markers turned out to be highly or fully informative, because they discriminate all four alleles of the biparental cross V3125 × ‘Börner’. One marker (GF01-60, derived from candidate SSR chr1_100137_2) was scored as monomorphic in this analysis, likely due to the small sequence difference of 2 bp in fragment size. In fact, exactly the 2 bp difference in fragment length were properly detected by Sanger sequencing, but are surely below the reliable resolution for a marker assay based on fragment analysis. The last of the 14 SSR markers, GF06-19 (SSR chr6_800_1), did not show any scoreable fragment for ‘Börner’, but amplified a single scoreable amplicon in V3125. Taken together, every method seems to have its limitations, but the different validation approaches demonstrated the high reliability and usability of our prediction approach for new SSR markers.

### Targeted mapping of the downy mildew resistance *Rpv14* by bulk segregant analysis

To assist the introgression of the downy mildew locus *Rpv14*, which has been mapped in ‘Börner’ to the lower arm of chr.5 (Ochssner et al., 2016), into *V. vinifera*, closely linked markers are required. For SNV marker-based association studies for *Rpv14*, 25 amplimers were designed targeting the lower arm of chr.5 of ‘Börner’, 17 of those turned out to be functional. As controls, two additional amplimers were designed, one at the top of chr.5 opposed to *Rpv14* and one on chr.14. All primers were designed based on BoeWGS1.0 contigs and SNV data (see Methods). The 76 amplicons obtained for 19 markers and four template DNAs (susceptible V3125, resistant donor ‘Börner’, the F1 bulks S and R) were sequenced and the trace files were subjected to peak area determination and evaluation.

All predicted SNVs within or between the BoeWGS1.0 contigs could be verified. In one case, we observed a tri-allelic SNV and removed the respective data points from further analyses. For SNVs from physically distinct contigs the peak ratios or SNV frequencies varied as initially expected for a bulked segregant analysis. By ordering BoeWGS1.0 contig pairs along the chr.5 of the reference genome sequence and allocating the corresponding SNVs or allele frequencies to these contigs the changes in the frequencies of the pools are obviously correlating with the candidate locus (Figure 2). For the control regions on chr.14 and at the top of chr.5 we hypothesized no selective pressure and therefore expected an allele frequency for the bulks of 75%. The observed frequencies on chr.14 fulfill the expectation. The frequencies in the north of chr.5 in the R bulk (55-60 %) differ to some extent from the expectation, where as in the S bulk the SNV frequency is as expected. This may be due to the low number of genotypes within the bulks. Even if only one out of the ten genotypes has no recombination between the target locus and the terminal part of the chromosome, the estimated frequency value changes theoretically by almost 10% of the expected value.

**Figure 2:**
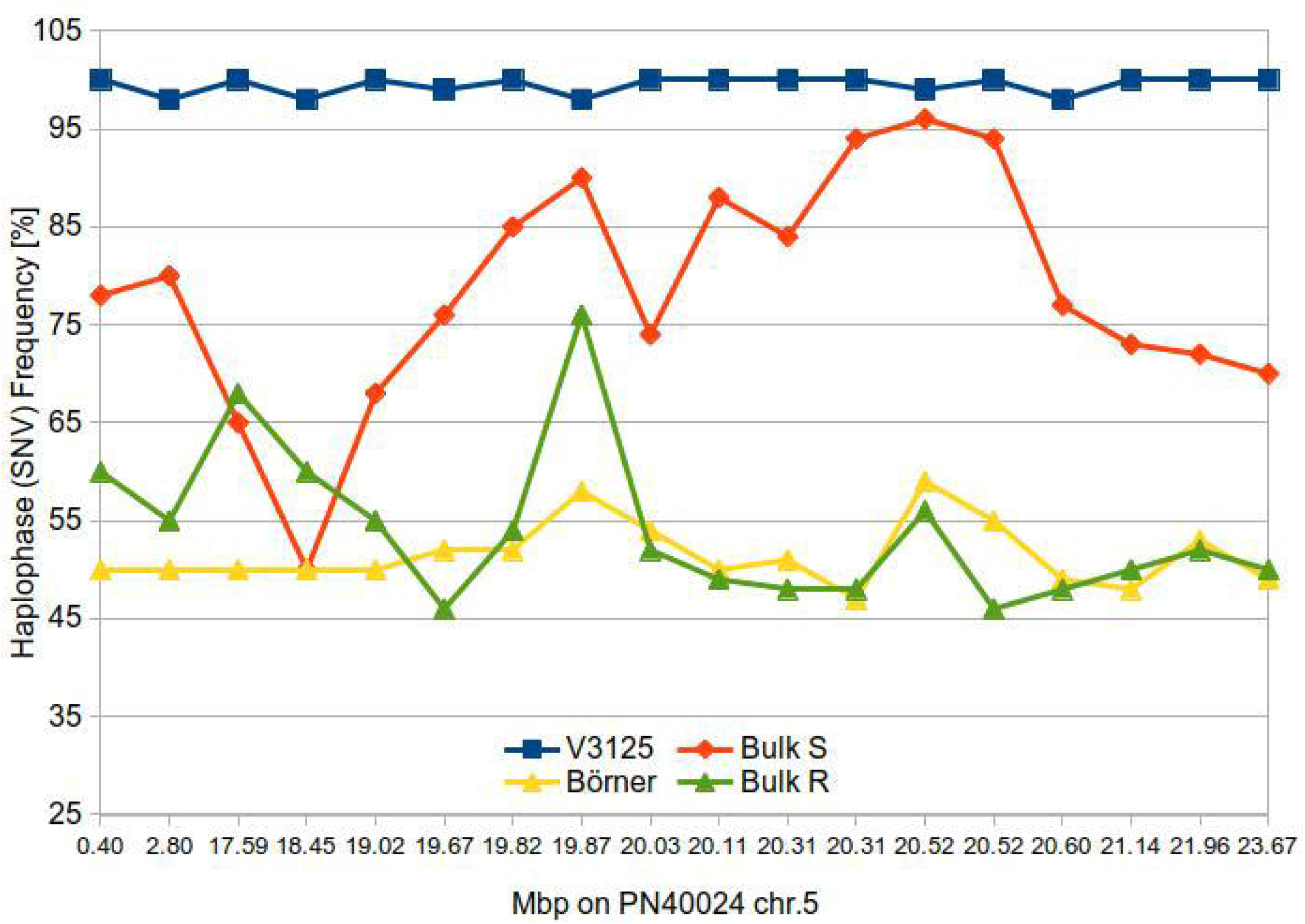
Haplotype frequency along the *Rpv14* target region of parental lines and bulks of susceptible and resistance F1 individuals.

Within the region of the candidate locus between approximately 19 and 21 Mbp, the resistant bulk has a nearly uniform estimated SNV frequency of 50%. This indicates that all resistant individuals in the bulk carry the same allele of the resistant ‘Börner’ parent (inherited from *V. cinerea*) (Ochssner et al., 2016). Since there are only a limited number of recombination events possible, the large interval is not surprising. At the same time, the SNV frequency in the susceptible bulk converges (continuously) to about 95%. Our results indicate that the variants of ‘Börner’, located between 19.67 Mbp and 20.6 Mbp in the corresponding reference sequence PN40024, are highly and specifically linked to the resistant trait. The center of the candidate locus with the highest linkage, showing SNV frequencies over 85% for the S bulk is an interval of 330 kbp between 20.31 and 20.6 Mbp. This is the region flanked by the ‘BoeWGS1.0’ contigs c35489, c244127 and c12059, c12060 (NCBI accessions CCJE01069526.1, CCJE01173868.1, CCJE01026433.1 and CCJE01045859.1).

## 4 Discussion

### Genome sequencing

For the development of molecular markers and to gain knowledge about genome regions as well as loci relevant for grapevine breeding we analyzed the genome of the resistant interspecific hybrid ‘Börner’ (*V. riparia* × *V. cinerea*) by whole genome shotgun (WGS) sequencing. The draft assembly BoeWGS1.0 covers a total of 618 Mbp, which corresponds to 62% of the expected 1 Gbp long diploid ‘Börner’ genome. Assessment by REAPR revealed some critical regions which could be caused by the incompletely resolved haplophases. Mapping of reads to at least in parts very similar sequences like e.g. separated alleles still poses a challenge. However, most of the contigs pass the read mapping-based assembly evaluation.

A major challenge during the *de novo* assembly of this dataset was to optimize the separation of sequences into contigs derived from both haplophases for subsequent generation of highly informative SSR and SNV marker assays. Different evaluations for the success of this separation result in somehow related, but in detail different numbers. BUSCO finds about 14% duplicated benchmarking genes, but since quite some genes of the reference set are not detected at all (BUSCO completeness is about 75%) this is probably a underestimation for the contig level since the relatively small contigs complicate gene calling. The total length of the assembly is about 120 Mbp longer than the haploid *Vitis* genome sequence, indicating that 240 of the 618 Mbp represent sequences from separated haplophases. The analyses of contig mappings against the reference genome sequence PN40024 indicates that 142 Mbp of the reference are covered by two BoeWGS1.0 contigs. Taken together, we assume that the BoeWGS1.0 assembly contains phase-separated sequence information for one quarter of the ‘Börner’ genome. Note that this information is not “phased throughout” since even if a contig pair is detected, we lack information about which of the two belongs to the *V. cineria* and which to the *V. riparia* haplophase. Anyway, since the regions for which separation worked are distributed throughout the genome, the data are a very good source for marker development.

Although the separation of sequences from both haplophases is great for molecular breeding applications and several genome-wide investigations, the short read derived BoeWGS1.0 assembly is still a draft genome sequence. The contig length still limits significantly downstream analyses like gene prediction or approaches to detect genomic rearrangements. The main reason for the fragmentation of the BoeWGS1.0 assembly are regions where the ancestor genomes of *V. cinerea* and *V. riparia* are quite similar. In fact, the frequency of SNVs and MNVs detected between the two haplophases of ‘Börner’ (i.e. *V. cinerea* and *V. riparia* haploid genomes) is in the same range as between ‘Börner’ and PN40024.

Currently, substantially longer read lengths (mean >20 kb) than those that were used here are provided by third generation single molecule sequencing technologies like those offered by Pacific Biosciences (PacBio) or Oxford Nanopore Technologies (ONT). The long reads can be used to solve the problem of bridging repeat sequences or transposable elements. The first *de novo* assembly of domesticated grapevine using PacBio sequencing was from *Vitis vinifera* cv. Cabernet Sauvingnon (Chin et al., 2016) followed recently by the diploid assembly of the cultivar Chardonnay (Roach et al., 2018; Zhou et al., 2019). The first and only wild *Vitis* genome sequence so far comes from the rootstock *Vitis riparia* ‘Gloire de Montpellier’ and reached considerable continuity by incoorperating PacBio and 10X Chromium (Girollet et al., 2019). However, assembling a highly heterozygous genotype, accurate phasing and aligning of *Vitis* haplotypes is still challenging. With regard to the data on hand and the complexity of this assembly, the application of dedicated scaffolding tools like scaffoldScaffolder (Bodily et al., 2015) or SSPACE (Boetzer et al., 2011) comes at a high risk of generating erroneous connections between contigs. Therefore, we calculated the BoeWGS1.0 assembly with very stringent parameters using CLC Genomics Workbench, which contains an implementation of a de Bruijn Graph assembler, and refrained from scaffolding.

In addition to the room for improvement for the assembly parameters, there is a significant fraction of the BoeWGS1.0 genome sequence currently not alignable and thus not assignable to the reference genome sequence. Likely, this genome fraction holds sequence information underlying important traits. Chromosomal anchoring of these contigs is a task that could be realized by the use of the provided SSR or SNV marker sequences of this study in an existing F1 mapping population (V3125 × ‘Börner’) (Fechter et al., 2014).

### Deduction of SSR sequences for marker development

The detected polymorphic SSR candidate loci are a very valuable resource for genome-wide marker development and genotyping. SSR markers are still the most abundant molecular marker type used for mapping in *Vitis* species and marker-assisted selection in grapevine breeding programs. Because of their very polymorphic nature SSRs often allow to clearly distinguish between more than two alleles. This is highly necessary when using F1 mapping populations derived from a cross of highly heterozygous parents, when following an allele in a phylogenetic tree or when identifying accessions (Cipriani et al., 2011). All these conditions fit to grapevine.

In this study, we were able to reliably predict thousands of candidates for SSR markers. Furthermore, we validated high prediction reliability, which is essentially limited only by design of specific primers, and demonstrated their applicability in *Vitis* genotyping approaches. The prediction schema for selection of promising SSR candidates that relied on aligning contigs representing separated haplophases instead of large amounts sequence reads turned out to be very successful. We were able to reliably detect, predict and select for longer sequence variations, caused by the variable number of repeated units. This is favorable for marker detection by fragment analysis, the currently mainly used method for SSR-based genotyping.

### Application of phased contig data for targeted mapping of the downy mildew resistance *Rpv14* of *Vitis*

For *V. riparia* (cv. Gloire de Montpellier) the mean distance between heterozygous SNVs is 217 bp (Girollet et al., 2019). For the heterozygous Pinot noir a frequency of 1 nucleotide variant per 100 bp and 1 InDel per 0.45 kb was described (Velasco et al., 2007). The overall variant frequency that we obtained between the ‘Börner’ genome sequence and the reference PN40024 was 1 variant per 68 bp. This number is likely to be a significant underestimation, because of the stringent read mapping parameters and the narrow confines for the coverage filter. Single SNVs or groups of nearby SNVs within the multiple alignments of two ‘Börner’ contig pairs and the PN40024 reference sequence were shown to be useful for the molecular identification and localization of a quantitative trait locus. The highly reliable SNVs detected in our study can function as molecular markers in a bulked segregant analysis as shown for *Rpv14* on chr.5 in our analysis. We were able to physically reduce the *Rpv14* target region to less than 500 kbp. In relation to the results from Ochßner et al. (Ochssner et al., 2016) the center of the target region is narrowed down from the north and the south with the center being still at 20.06 Mbp.

Further research should focus on the identification of F1 individuals with additional recombinations in the target region. Subsequently, the work presented here allows quick access to additional markers which can be used to determine the recombination points that then can be correlated with resistance genes from candidate gene predictions.

## Author Contributions

DH and LH conceived and planned the experiments. DH, PV, LH designed and performed the experiments. TR, DH, BP calculated the data. RT, BW, DH conceived the original idea and supervised the project. DH, RT and BW wrote the manuscript with input from all authors. All authors interpreted and discussed results.

## Acknowledgments

We thank Willy Keller, Andreas Preiß, and Andreas Melnik for the excellent technical support. The authors also wish to thank the members of the Chair of Genetics and Genomics of Plants for their support. We gratefully acknowledge the financial support from BMBF through Projektträger Jülich. We also acknowledge support for the Article Processing Charge by the Deutsche Forschungsgemeinschaft and the Open Access Publication Fund of Bielefeld University.

## References

Adam-Blondon, A.-F., Jaillon, O., Vezzulli, S., Zharkikh, A., Troggio, M., and Velasco, R. (2011). “Genome Sequence Initiatives,” in Genetics, Genomics and Breeding of Grapes, eds. A.-F. Adam-Blondon, J.M. Martinez-Zapater & C. Kole.).

Alkan, C., Sajjadian, S., and Eichler, E.E. (2011). Limitations of next-generation genome sequence assembly. Nature Methods 8(1), 61–65.

Altschul, S.F., Gish, W., Miller, W., Myers, E.W., and Lipman, D.J. (1990). Basic local alignment search tool. Journal of Molecular Biology 215(3), 403–410.

Barba, P., Cadle-Davidson, L., Harriman, J., Glaubitz, J.C., Brooks, S., Hyma, K., et al. (2014). Grapevine powdery mildew resistance and susceptibility loci identified on a high-resolution SNP map. Theoretical and Applied Genetics 127(1), 73–84.

Barcellos, L.F., Klitz, W., Field, L.L., Tobias, R., Bowcock, A.M., Wilson, R., et al. (1997). Association mapping of disease loci, by use of a pooled DNA genomic screen. American Journal of Human Genetics 61(3), 734–747.

Bellin, D., Peressotti, E., Merdinoglu, D., Wiedemann-Merdinoglu, S., Adam-Blondon, A.F., Cipriani, G., et al. (2009). Resistance to Plasmopara viticola in grapevine ‘Bianca’ is controlled by a major dominant gene causing localised necrosis at the infection site. Theoretical and Applied Genetics 120, 1.

Bodily, P.M., Fujimoto, M., Ortega, C., Okuda, N., Price, J.C., Clement, M.J., et al. (2015). Heterozygous genome assembly via binary classification of homologous sequence. BMC Bioinformatics 16 Suppl 7(S5), 1471-2105-1416-S1477-S1475.

Boetzer, M., Henkel, C.V., Jansen, H.J., Butler, D., and Pirovano, W. (2011). Scaffolding pre-assembled contigs using SSPACE. Bioinformatics 27(4), 578–579.

Bolger, A.M., Lohse, M., and Usadel, B. (2014). Trimmomatic: a flexible trimmer for Illumina sequence data. Bioinformatics 30(15), 2114–2120.

Burton, J.N., Adey, A., Patwardhan, R.P., Qiu, R., Kitzman, J.O., and Shendure, J. (2013). Chromosome-scale scaffolding of de novo genome assemblies based on chromatin interactions. Nature Biotechnology 31(12), 1119–1125.

Camacho, C., Coulouris, G., Avagyan, V., Ma, N., Papadopoulos, J., Bealer, K., et al. (2009). BLAST+: architecture and applications. BMC Bioinformatics 10, 421.

Capistrano-Gossmann, G.G., Ries, D., Holtgräwe, D., Minoche, A., Kraft, T., Frerichmann, S.L.M., et al. (2017). Crop wild relative populations of Beta vulgaris allow direct mapping of agronomically important genes. Nature Communications 8(15708).

Carr, I.M., Robinson, J.I., Dimitriou, R., Markham, A.F., Morgan, A.W., and Bonthron, D.T. (2009). Inferring relative proportions of DNA variants from sequencing electropherograms. Bioinformatics 25(24), 3244–3250.

Chin, C.S., Peluso, P., Sedlazeck, F.J., Nattestad, M., Concepcion, G.T., Clum, A., et al. (2016). Phased diploid genome assembly with single-molecule real-time sequencing. Nature Methods 13(12), 1050–1054.

Cipriani, G., Di Gaspero, G., Canaguier, A., Jusseaume, J., Tassin, J., Lemainque, A., et al. (2011). “Molecular Linkage Maps: Strategies, Resources and Achievements,” in Genetics, Genomics and Breeding of Grapes, eds. A.-F. Adam-Blondon, J.M. Martinez-Zapater & C. Kole.).

Di Genova, A., Almeida, A.M., Muñoz-Espinoza, C., Vizoso, P., Travisany, D., Moraga, C., et al. (2014). Whole genome comparison between table and wine grapes reveals a comprehensive catalog of structural variants. BMC Plant Biology 14(7).

Dohm, J.C., Lange, C., Holtgräwe, D., Sörensen, T.R., Borchardt, D., Schulz, B., et al. (2012). Palaeohexaploid ancestry for Caryophyllales inferred from extensive gene-based physical and genetic mapping of the sugar beet genome (Beta vulgaris). The Plant Journal 70(3), 528–540.

Fechter, I., Hausmann, L., Zyprian, E., Daum, M., Holtgräwe, D., Weisshaar, B., et al. (2014). QTL analysis of flowering time and ripening traits suggests an impact of a genomic region on linkage group 1 in Vitis. Theoretical and Applied Genetics 127(9), 1857–1872.

Girollet, N., Rubio, B., and Bert, P.F. (2019). De novo phased assembly of the Vitis riparia grape genome. Scientific Data 6(1), 127.

Hausmann, L., Maul, E., Ganesch, A., and Töpfer, R. (2019). Overview of genetic loci for traits in grapevine and their integration into the VIVC database. Acta Hortic 1248, 2019.1248.2032.

Herzog, E., Töpfer, R., Hausmann, L., Eibach, R., and Frisch, M. (2013). Selection strategies for marker-assisted background selection with chromosome-wise SSR multiplexes in pseudo-backcross programs for grapevine breeding. Vitis 52(4), 193–196.

Hunt, M., Kikuchi, T., Sanders, M., Newbold, C., Berriman, M., and Otto, T.D. (2013). REAPR: a universal tool for genome assembly evaluation. Genome Biology 14(5), R47.

Hyma, K.E., Barba, P., Wang, M., Londo, J.P., Acharya, C.B., Mitchell, S.E., et al. (2015). Heterozygous Mapping Strategy (HetMappS) for High Resolution Genotyping-By-Sequencing Markers: A Case Study in Grapevine. PLoS ONE 10(8).

Jaillon, O., Aury, J.M., Noel, B., Policriti, A., Clepet, C., Casagrande, A., et al. (2007). The grapevine genome sequence suggests ancestral hexaploidization in major angiosperm phyla. Nature 449(7161), 463–467. doi: 10.1038/nature06148.

Kent, W.J. (2002). BLAT--the BLAST-like alignment tool. Genome Research 12(4), 656–664.

Li, C., Lin, F., An, D., Wang, W., and Huang, R. (2018). Genome Sequencing and Assembly by Long Reads in Plants. Genes 9(1).

Li, H. (2013). Aligning sequence reads, clone sequences and assembly contigs with BWA-MEM. arXiv,1303.3997v1302 (Preprint posted May 1326, 2013).

Lodhi, M.A., and Reisch, B.I. (1995). Nuclear DNA content of Vitis species, cultivars, and other genera of the Vitaceae. Theoretical and Applied Genetics 90(1), 11–16.

Merdinoglu, D., Wiedemann-Merdinoglu, S., Coste, P., Dumas, V., Haetty, S., Butterlin, G., et al. (2003). Genetic Analysis of Downy Mildew Resistance Derived from Muscadinia rotundifolia. Acta Hort 603.

Michael, T.P., and Jackson, S. (2013). The First 50 Plant Genomes. The Plant Genome 6(2).

Michelmore, R.W., Paran, I., and Kesseli, R.V. (1991). Identification of markers linked to disease-resistance genes by bulked segregant analysis: a rapid method to detect markers in specific genomic regions by using segregating populations. Proceedings of the National Academy of Sciences of the United States of America 88(21), 9828–9832.

Ochssner, I., Haussmann, L., and Töpfer, R. (2016). Rpv14, a new genetic source for Plasmopara viticola resistance conferred by vitis cinerea. Vitis 55, 79–81.

Rex, F., Fechter, I., Hausmann, L., and Töpfer, R. (2014). QTL mapping of black rot (Guignardia bidwellii) resistance in the grapevine rootstock ‘Börner’ (V. riparia Gm183 × V. cinerea Arnold). Theoretical and Applied Genetics 127(7), 1667–1677.

Roach, M.J., Johnson, D.L., Bohlmann, J., van Vuuren, H.J.J., Jones, S.J.M., Pretorius, I.S., et al. (2018). Population sequencing reveals clonal diversity and ancestral inbreeding in the grapevine cultivar Chardonnay. PLoS Genetics 14(11), e1007807.

Schwander, F., Eibach, R., Fechter, I., Hausmann, L., Zyprian, E., and Töpfer, R. (2012). Rpv10: a new locus from the Asian Vitis gene pool for pyramiding downy mildew resistance loci in grapevine. Theoretical and Applied Genetics 124(1), 163–176.

Sham, P., Bader, J.S., Craig, I., O’Donovan, M., and Owen, M. (2002). DNA Pooling: a tool for large-scale association studies. Nature Reviews Genetics 3(11), 862–871.

Shaw, S.H., Carrasquillo, M.M., Kashuk, C., Puffenberger, E.G., and Chakravarti, A. (1998). Allele frequency distributions in pooled DNA samples: applications to mapping complex disease genes. Genome Research 8(2), 111–123.

Simão, F.A., Waterhouse, R.M., Ioannidis, P., Kriventseva, E.V., and Zdobnov, E.M. (2015). BUSCO: assessing genome assembly and annotation completeness with single-copy orthologs. Bioinformatics 31(19), 3210–3212.

Taheri, S., Lee Abdullah, T., Yusop, M.R., Hanafi, M.M., Sahebi, M., Azizi, P., et al. (2018). Mining and Development of Novel SSR Markers Using Next Generation Sequencing (NGS) Data in Plants. Molecules 23(2), E399.

Thiel, T., Michalek, W., Varshney, R.K., and Graner, A. (2003). Exploiting EST databases for the development and characterization of gene-derived SSR-markers in barley (Hordeum vulgare L.). Theoretical and Applied Genetics 106(3), 411–422.

Töpfer, R., and Eibach, R. (2016). Breeding the next-generation disease-resistant grapevine varieties. Wine & Viticulture Journal.

Untergasser, A., Cutcutache, I., Koressaar, T., Ye, J., Faircloth, B.C., Remm, M., et al. (2012). Primer3--new capabilities and interfaces. Nucleic Acids Research 40(15).

VanBuren, R., Bryant, D., Edger, P.P., Tang, H., Burgess, D., Challabathula, D., et al. (2015). Single-molecule sequencing of the desiccation-tolerant grass Oropetium thomaeum. Nature 527(7579), 508–511.

Velasco, R., Zharkikh, A., Troggio, M., Cartwright, D.A., Cestaro, A., Pruss, D., et al. (2007). A high quality draft consensus sequence of the genome of a heterozygous grapevine variety. PLoS One 2(12).

Venuti, S., Copetti, D., Foria, S., Falginella, L., Hoffmann, S., Bellin, D., et al. (2013). Historical introgression of the downy mildew resistance gene Rpv12 from the Asian species Vitis amurensis into grapevine varieties. PLoS One 8(4), e61228.

Welter, L.J., Göktürk-Baydar, N., Akkurt, M., Maul, E., Eibach, R., Töpfer, R., et al. (2007). Genetic mapping and localization of quantitative trait loci affecting fungal disease resistance and leaf morphology in grapevine (Vitis vinifera L). Molecular Breeding 20, 359–374.

Zhang, J., Hausmann, L., Eibach, R., Welter, L.J., Töpfer, R., and Zyprian, E.M. (2009). A framework map from grapevine V3125 (Vitis vinifera ‘Schiava grossa’ × ‘Riesling’) × rootstock cultivar ‘Börner’ (Vitis riparia x Vitis cinerea) to localize genetic determinants of phylloxera root resistance. Theoretical and Applied Genetics 119(6), 1039–1051.

Zhou, Y., Minio, A., Massonnet, M., Solares, E., Lv, Y., Beridze, T., et al. (2019). The population genetics of structural variants in grapevine domestication. Nature Plants 5(9), 965–979.

Zimmer, R., and Verrinder Gibbins, A.M. (1997). Construction and characterization of a large-fragment chicken bacterial artificial chromosome library. Genomics 42(2), 217–226.

