## Supplementary figures and images for "A Partially Phase-Separated Genome Sequence Assembly of the *Vitis* Rootstock ‘Börner’ (*Vitis riparia* x *Vitis cinerea*) and its Exploitation for Marker Development and Targeted Mapping"

### Supplementary Figure 1

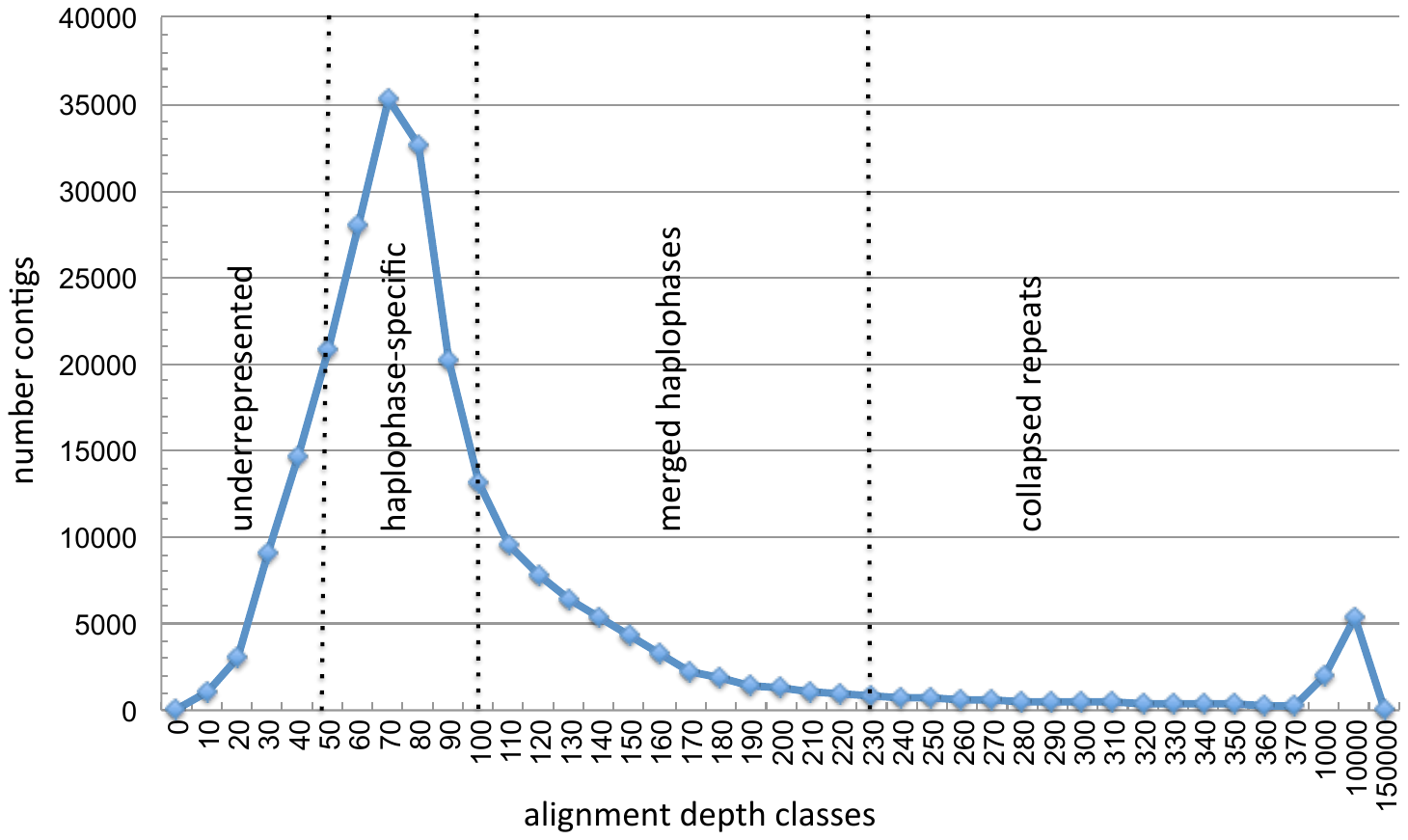

**Supplementary Figure 1**. Allele depth classes of BoeWGS1.0 contigs.
